# Human embryonic tanycyte: heterogeneity and developmental trajectory

**DOI:** 10.1101/2023.01.03.522431

**Authors:** Yuan Li

## Abstract

Disruption of energy homeostasis may cause diseases such as obesity and diabetes that affect millions of people every year. The adult hypothalamic stem cells, tanycytes, play critical roles in helping hypothalamic neurons maintain energy homeostasis, however the developmental trajectory of tanycytes especially in human still awaits to be discovered. In the current study, we for the first time use human embryonic single cell transcriptomics data to distinguish RAX^+^ tanycytes from RAX^+^ neural progenitors, explore human embryonic tanycyte heterogeneity, and unravel their developing trajectories. We found human embryonic tanycytes share similar subtypes with adult rodent tanycytes (α and β). We also discovered that radial glia markers *FABP7* as well as astrocyte marker (e.g. *AQP4*) etc, are characteristics of tanycytes that distinguish them from RAX^+^ neural progenitors, and the α and β tanycytes follow different developmental trajectories. Our study represents a pioneer work on human embryonic tanycytes.

## Introduction

Disruption of energy homeostasis may cause diseases such as obesity and diabetes that affect millions of people every year, compromising their quality of life, even risking their life. Tight regulation of energy homeostasis is controlled by central nervous system, especially the hypothalamus that functions as a hub of a complex neural network that precisely coordinates energy expenditure, food intake and blood sugar levels (Bolborea and Langlet 2021, Langlet 2019, Luquet and Magnan 2009). The ependymoglial cell, tanycyte has recently been shown to play critical roles in passing the nutrient sufficiency information carried by circulating signals (such as leptin, ghrelin and insulin) to different hypothalamus neurons, such as the orexigenic agouti-related peptide (AGRP) and the anorexigenic proopiomelanocortin (POMC) neurons (Porniece Kumar et al. 2021, Duquenne et al. 2021, Uriarte Donati et al. 2019). The function of tanycytes largely benefit from their unique location as the floor and side wall of at least part of the 3rd ventricle, where they have their apical side in direct contact with the cerebrospinal fluid (CSF) from the ventricle, and send their long processes to the fenestrated capillaries in the median eminence (ME) and to hypothalamus neurons (Prevot et al. 2018). Furthermore, at least some tanycytes have been shown to also function as adult neural progenitors that can give rise to neurons and astrocyte, contributing to neural plasticity (Prevot et al. 2018). Tanycyte is a heterogeneous cell types, comprising two α subtypes and two β subtypes in adult rodent hypothalamus (Campbell et al. 2017, Sullivan et al. 2022). Those tanycyte subtypes occupy different areas along the floor (β) and lateral wall (α) of the third ventricle (Campbell et al. 2017). Both α and β tanycytes are characterised by their expression of progenitor markers RAX and VIM, as well as factors involved in energy balance (FGF10) and thyroid hormone regulation (DIO2) (Campbell et al. 2017, Sullivan et al. 2022). While rodent α tanycytes specifically express NR2E1, FABP7 and VCAN, rodent β tanycytes show a high expression of COL25A1, ADM, as well as FGF1R that is important in glucose homeostasis (Campbell et al. 2017, Bentsen et al. 2020).

Regarding the developmental trajectory of tanycytes, Chinnaiya et al. (2022) have identified a tuberal hypothalamus progenitor population that shares very similar gene expression profile with adult rodent tanycytes: they both express RAX, VIM and FGF10. Chinnaiya et al. (2022) collected single cell transcriptomics data for those cells from a chicken dataset (Kim et al. 2022) and compared them between different embryonic stages, they found that as development moved forward, radial glia markers VIM and FABP7 got upregulated in embryonic tanycyte cells, likely reflecting a transition from neural progenitors towards radial glia-like tanycytes (Chinnaiya et al. 2022). However, how this developmental process looks like in human still remains to be discovered. In the current study, we have collected and analysed RAX^+^ progenitor clusters from a human embryonic hypothalamus dataset, clustered them, mapped them onto a mouse reference data set, performed trajectories analysis and did machine learning based identification of dynamically regulated genes, with the aim to unveil their transcriptomic diversity and sort out their developmental trajectories.

## Materials and Methods

### Gather potential embryonic tanycytes

A human hypothalamus single cell transcriptomics dataset has been downloaded from the NCBI Gene Expression Omnibus (GEO) database (accession number: GSE169109). The dataset includes Gestational week/GW 7, 8, 10, 12, 15, 18 and 20, and each developmental stage (except GW8 that had low-quality data) was processed separately with Seurat v.4.1.0 (Satija et al. 2015). To acquire potential embryonic tanycyte, we collected all RAX^+^VIM^+^STMN2^low^ progenitor clusters from each developmental stages, and reprocessed them together, the newly selected dataset was then clustered using Seurat v.4.1.0. with default setting, from which several clusters appeared to be high in STMN2 expression and/or low in RAX and were removed from further analyses (Figure S1_A, B). For the remained 1947 RAX^+^ progenitor cells, cell-wise cell cycle scores were calculated based on a predefined list of cell cycle genes (https://github.com/scverse/scanpy_usage/blob/master/180209_cell_cycle/data/regev_lab_cell_cycle_genes.txt). Tanycytes normally have low proliferative activity (Prevot et al. 2018), and we observed that most of the non-cycling cells (in G1 phase) were from later developmental stages (GW15/18/20) (Fig S1_C) when the progenitors have developed more towards the tanycyte fate; only non-cycling cells were maintained for further analyses (Fig S1_D).

### Label transfer to identify tanycytes

To check exactly which of the gathered RAX+ progenitor clusters are embryonic tanycytes, we have first reclustered the non-cycling RAX+ progenitors and then, on which we performed MapQuery analysis as implemented in Seurat v.4.1.0, the reference datasets used for this analysis was from mouse hypothalamus (Campbell et al. 2017), where the tanycyte populations were clearly identified and characterized together with neurons, astrocytes, ependymocytes, oligodendrocytes, VLMCs, endothelial cells, microglia and pars tuberalis. At louvain clustering resolution 1.5, in total 13 clusters were acquired (Figure 1_B), and six of which were mainly mapped onto the reference neurons, suggesting a neuronal progenitor identity of those clusters (Table S1). While all the rest 7 (except cluster Nr. 1) were largely mapped onto reference tanycyte (Table S1). Louvain cluster Nr. 1 was primarily predicted to be pars tuberalis, which is part of the pituitary gland and is characterized by a high expression of CCK and PITX2 (Campbell et al. 2017). However, neither of these two genes were highly/specifically expressed in cluster Nr. 1, instead, it was the tanycyte marker CRYM and FRZB (Campbell et al. 2017) that had strong expression in this cluster (Figure S2_A), therefore, we considered this cluster to be another embryonic tanycyte population (Figure 1_B).

**Figure 1.**
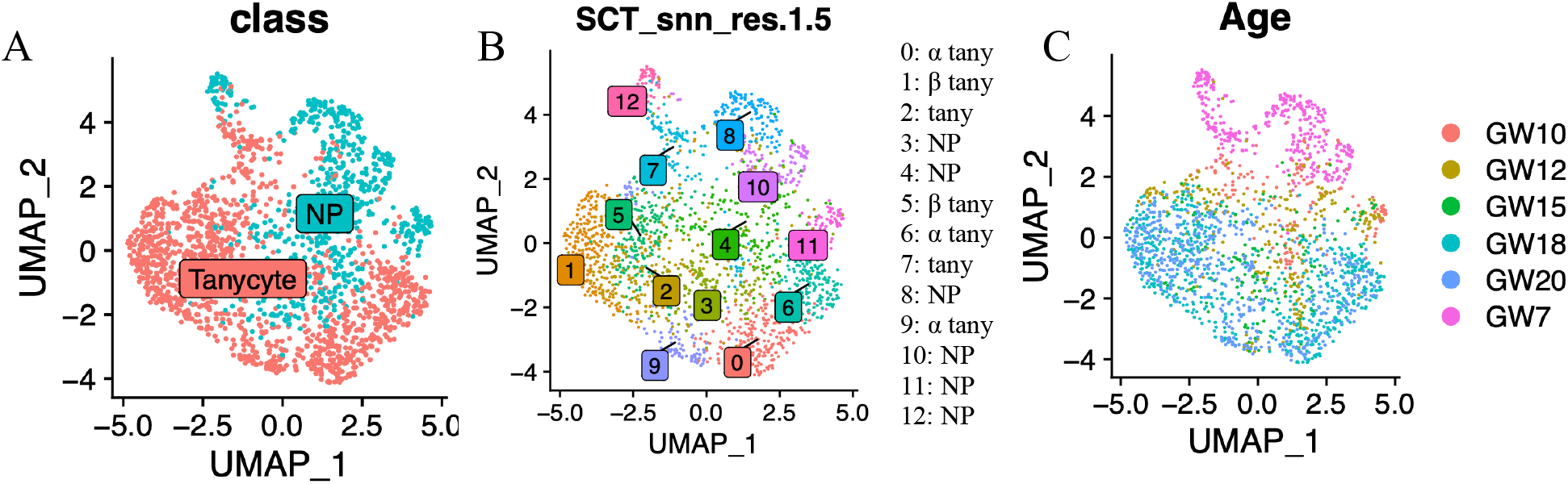
RAX^+^ progenitor heterogeneity. (A) UMAP plot of 1947 cells colored based on cell type category (neural progenitor/NP or tanycyte). (B) UMAP plot showing the results of Louvain clustering at a resolution of 1.5 as well as the predicted cell type for each cluster. (C) UMAP plot of the same cells labeled according to their embryonic stages (GW: gestational week).

### Comparison between RAX+ tanycyte and RAX+ neural progenitors

Most of the predicted neural progenitor clusters are from early developmental stages, (GW7/10/12/15), while the embryonic tanycyte cells were mostly found at later developmental stages (GW18 and GW10, Figure 1_C). To validate the Seurat label prediction result, we have grouped the neural progenitor clusters and the tanycyte lineage clusters together respectively (Figure 1_A), and identified differentially expressed genes between the two groups with Seurat v.4.1.0.

### Trajectory inference and machine learning based identification of lineage-wise dynamically regulated genes

To infer cell lineages and chart their developmental progression from progenitors towards different tanycyte subtypes, we subset the tanycyte clusters and performed a cell trajectory analysis on them using the R package Slingshot (Street et al. 2018). Slingshot inference of cell lineages was based on the Louvain clusters at a clustering resolution of 0.5 (root: cluster nr 5, most cells of which are from the earliest time point GW7), where each cluster was considered as a distinct cellular state. A bifurcating lineage structure that connects those cellular stages was constructed as a Minimum Spanning Tree (MST) (Tewarie et al. 2015) from cells that was organized in a UMAP-created reduced-dimensional space. From this global lineage structure, the developmental progression (pseudotime) for each cell was estimated, lineage by lineage, using the Simultaneous Principal Curves method (Hastie et al. 1989).

To identify dynamic temporal genes along the two tanycyte developmental lineages identified with Slingshot, we have built a random forest regression model using the R package tidymodels v.0.1.4.9000 (Kuhn & Wickham, 2020). The count matrix used for model construction was from the counts slot of the SCT assay, including top 2000 highly variable genes (HVGs), and then scaled using the R function scale (Becker et al. 1988). This preprocessed data matrix was next split into a training (3/4 of the data) and a testing dataset, and based on the training dataset, 1400 decision trees were constructed. For each tree, the dataset was split into subsets recursively until the subsets reached the minimum cell number, and every split was made in a way that the data impurity in the parent datasets was reduced the most, and data impurity was evaluated based on selected HVGs. Here, the minimum cell number per subset and the selected HVG number at each split were acquired based on model tuning via grid search. The prediction performance of the constructed model (decision tree ensemble) was evaluated on the testing dataset, where the mean of the predicted pseudotime from the 1400 decision trees was compared with the known pseudotime for each cell. As a result, we got a RMSE (root mean squared error) value of 1.70/1.51 and a MAE (mean absolute deviation) value of 1.15/1.07 respectively for the α and β tanycyte lineages. From the constructed random forest model, top 50 genes (plus PENK and VCAN for the α tanycyte lineage) regarding their importance in the pseudotime prediction were selected as the dynamic temporal genes (Figure S), and the gene expression heatmap of those temporal genes along the lineage trajectory were drawn with the R function Heatmap as implemented within ComplexHeatmap v.2.6.2 (Gu 2022).

## Results

### RAX+ tanycytes vs. RAX+ neural progenitors

Differentially expressed gene analysis shows that tanycyte marker genes CRYM, MT2A and FRZB (Campbell et al. 2017) were unregulated in embryonic tanycytes, while anterior tuberal hypothalamus progenitor markers HES5 and SHH (Chinnaiya et al. 2022) were upregulated in the neural progenitor clusters (Figure 2_A, Table S2). Apart from that, astrocyte markers SLC1A3/GLAST, FABP7/BLBP, GFAP, AQP4 and S100B were also expressed more in the tanycyte clusters comparing to the neural progenitor clusters (Figures 2 and 3_B).

**Figure 2.**
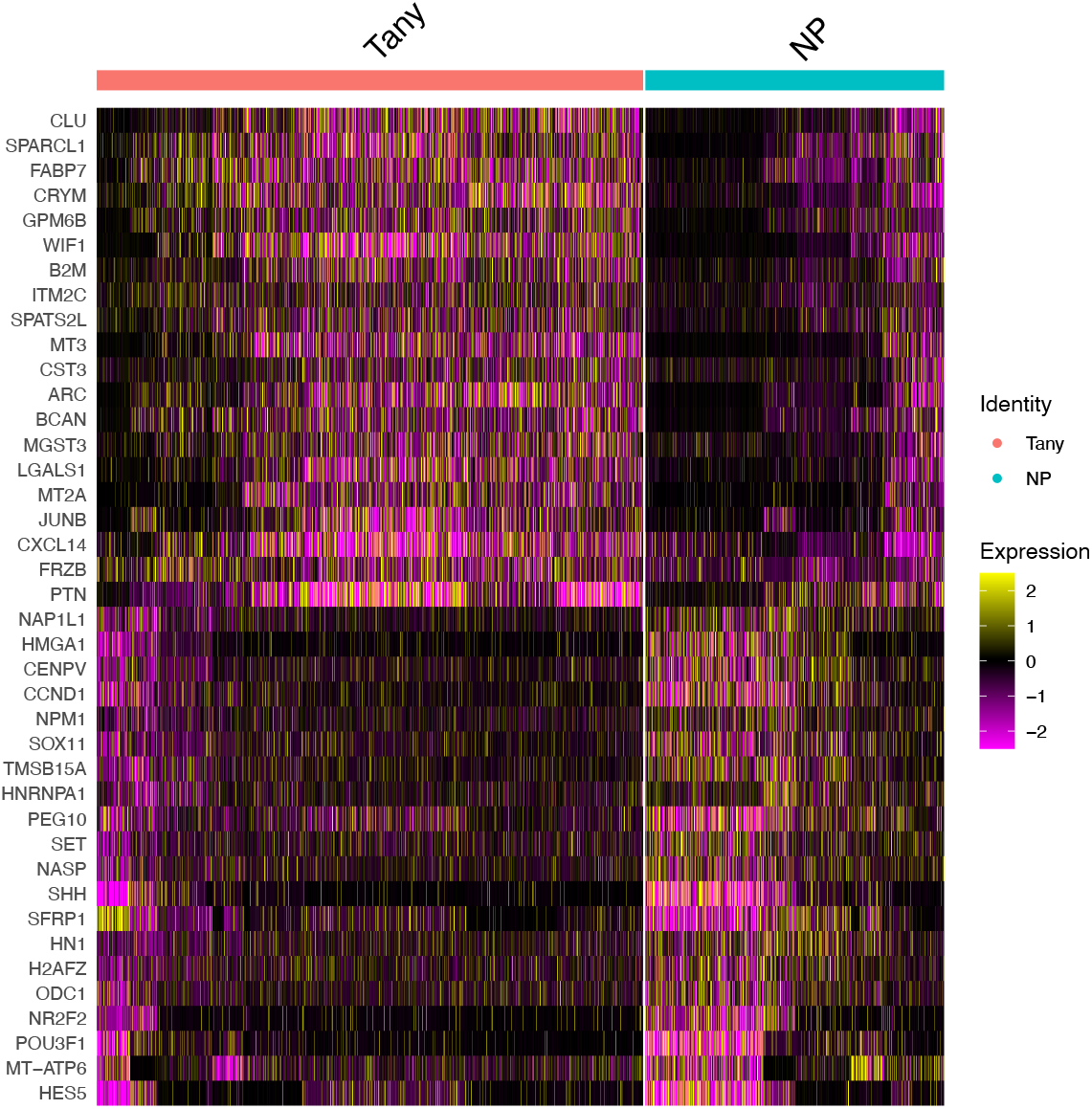
RAX^+^ embryonic tanycytes vs RAX^+^ neural progenitors. Gene expression heatmap of differentially expressed genes between RAX^+^ embryonic tanycytes and RAX^+^ neural progenitors.

**Figure 3.**
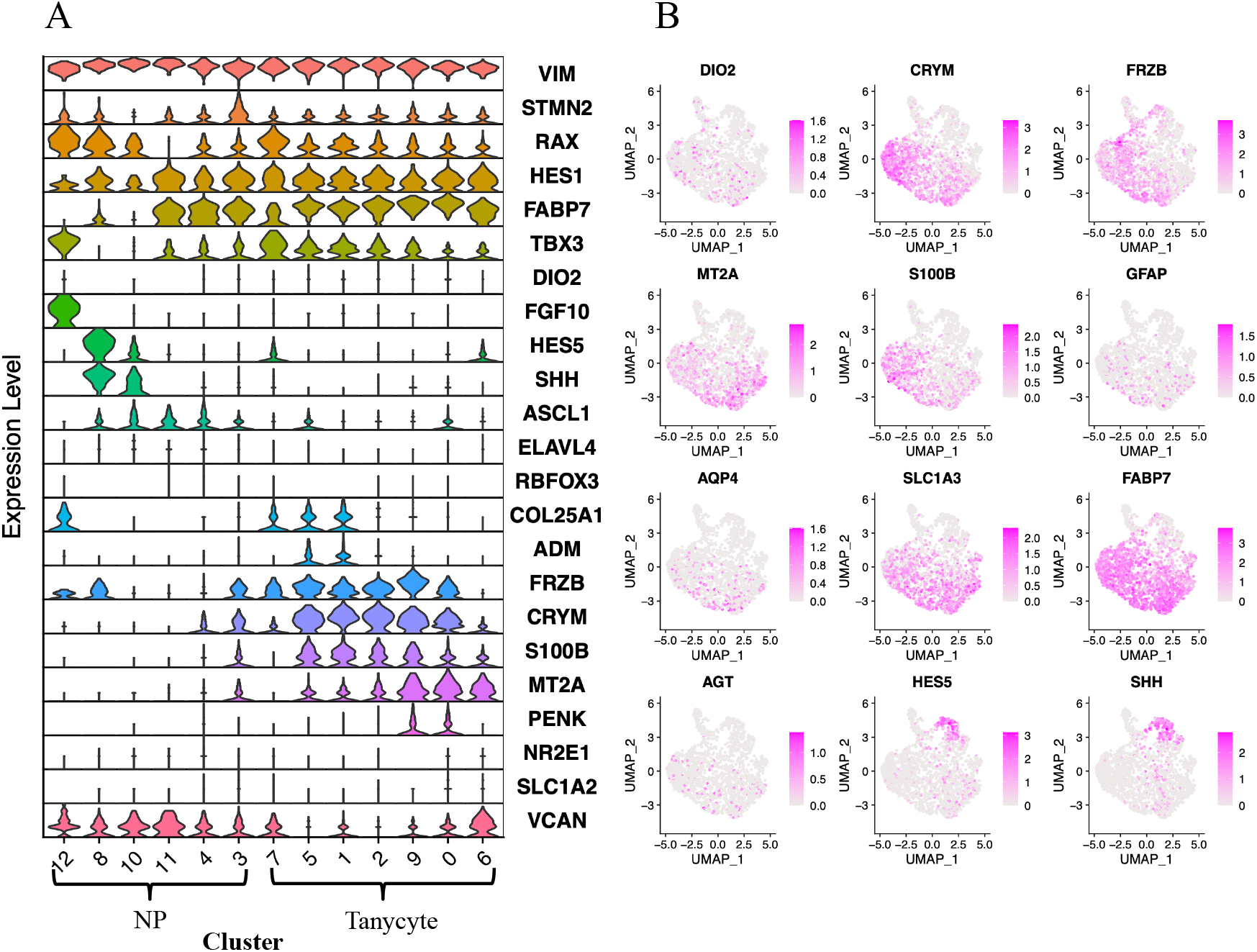
RAX^+^ embryonic tanycyte heterogeneity. (A) Stacked violin plot showing the expression levels of a list of marker genes within the 13 clusters from Figure 1_B. (B) UMAP plots showing cell-wise expression levels of a list of marker genes.

### Embryonic α and β tanycytes

A close examination of the tanycyte clusters reveal clear heterogeneity, which can be approximately grouped into two subgroups: in one subgroup, clusters 5 and 1 strongly express ADM and COL25A1, corresponding to the mouse β tanycyte, while in the second subgroup, clusters 9, 0 and 6 show specific expression of MT2A, NR2E1, SLC1A2 or VCAN, corresponding to mouse α tanycyte (Figure 3). Clusters 2 appears to be intermediate between the α and β tanycyte subgroups, and expresses markers that are shared by both, such as FRZB and CRYM (Figure 3). Cluster 7 occurs at the earliest developmental stage (mostly at GW7 and GW10) (Figure 1).

### Trajectory analyses revealing different developmental trajectories for the α and β tanycytes

Slingshot trajectory analysis has identified two developmental lineages among the tanycyte cells, one gives rise to β tanycyte, while the second one leads to the α tanycyte (Figure 4_A). A list of dynamically regulated genes were identified during each of the differentiation processes, a pseudotime ordering of the top 50 dynamically regulated genes along the β tanycyte developmental lineage unravel that, cell positive for *SFRP1* were present at an early stage of the developmental trajectory. SFRP1 is a Wnt pathway antagonist representing a possible signal for promoting quiescence of progenitors at the aged human subventricular zone (Donegal et al. 2022). Shortly after the start, *NNAT* and the subsequent *FABP7* got unregulated. NNAT plays a role in metabolic regulation and it has been shown to be expressed by tanycyte not by neurons in the arcuate nucleus (Vrang et al. 2010), *FABP7* is a radial glia marker (De Rosa et al. 2012). The upregulation of both genes may reflect the transition from a progenitor state towards a radial glia-like tanycyte fate (Chinnaiya et al. 2022). The uplift of *FABP7* is first followed by the rise of *CLU* and *APOE*, then by the increased expression of a marker gene *CRYM* shared by both α and β tanycytes (Campbell et al. 2017). APOE is involved in fat metabolism in mammal (Stolerman 2010), while CLU is a molecular chaperone that aids protein folding (Koltai 2014). At the end of the β tanycyte developmental trajectory, we got enhanced expression of *FRZB* (a more β tanycyte marker) (Campbell et al. 2017). An outstanding phenomenon of the β tanycyte developmental trajectory is the strong expression of *CXCL14* at later developmental stages.

**Figure 4.**
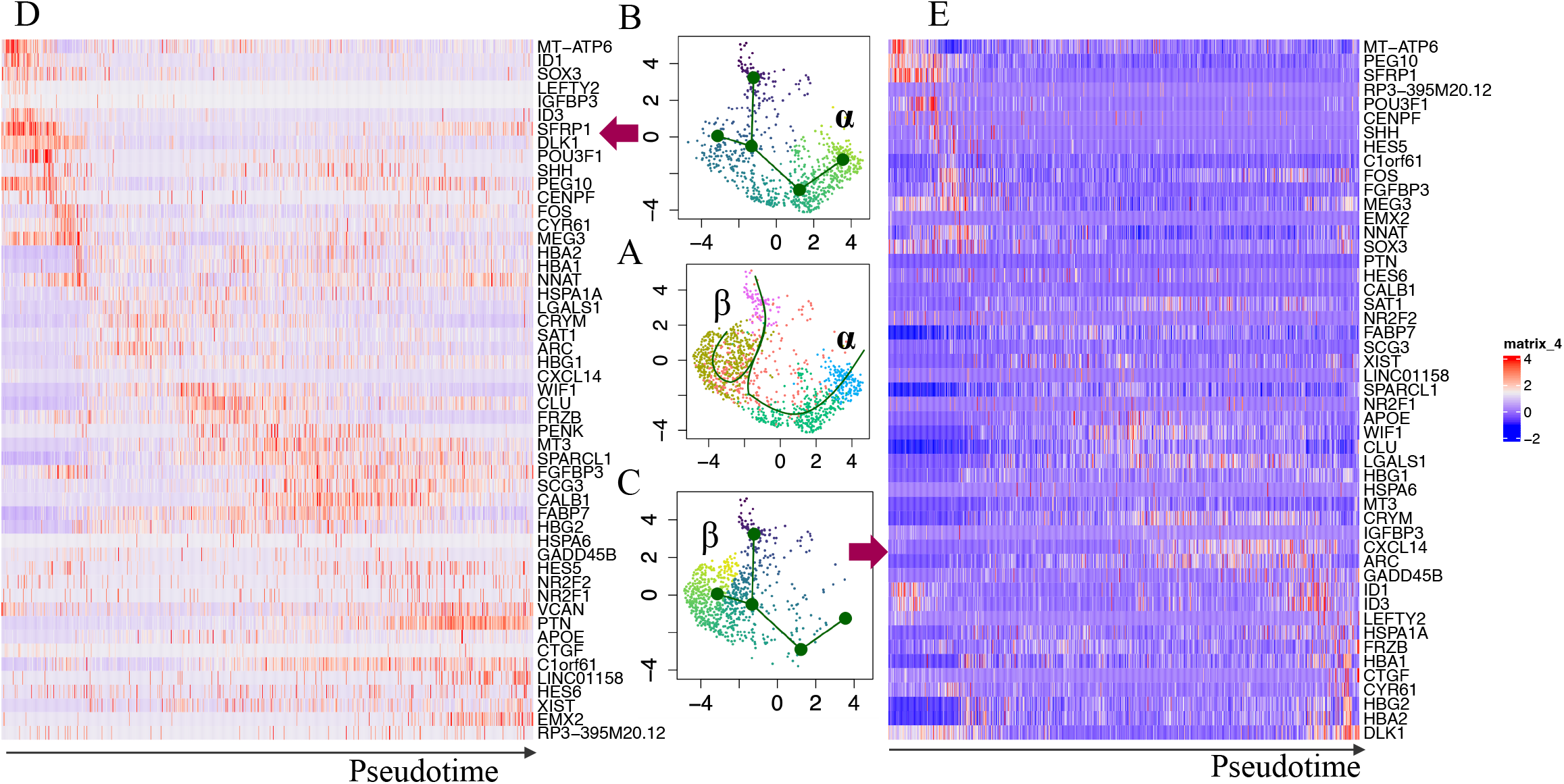
Trajectory analyses revealing different developmental trajectories for the α and β tanycyte. (A) UMAP showing the two tanycyte lineages. Cells are colored according to the Louvain clusters acquired at a resolution of 0.5. (B) & (C) UMAP plots showing, respectively, the α and β developmental trajectory, with cell colored based on pseudotime (root: dark blue). (D) & (E) heatmaps of the expression levels of dynamically regulated genes along, respectively, the α and β developmental trajectory.

The developmental trajectory of α tanycytes shared the same origin as for β tanycyte, and it also expresses *SFRP1*, as well as *ID1, ID3* and *LEFTY2*. Those progenitor gene expression enhancement is followed by the uplift of DLK1, POU3F1 and later on the *NNAT* genes. Afterwards, *FABP7*, the tanycyte markers *CRYM, FRZB* and *PENK*, as well as *CLU* are upregulated at different intermediate developmental stages. Towards the end of the developmental trajectory, the expression of the α tanycyte marker *VCAN* is enhanced. Similar to the *CXCL14* gene for β tanycyte, *PTN* shows an outstandingly high gene expression towards the end of the α tanycyte developmental trajectory.

## Discussion

The adult hypothalamic stem cells, tanycytes, play critical roles in helping hypothalamus neurons maintain metabolic homeostasis in mammals. In the current study, we for the first time use human embryonic single cell transcriptomics data to distinguish RAX+ tanycytes from RAX+ neural progenitors, explore tanycyte heterogeneity, and unravel their developing trajectories. The exploration of embryonic tanycytes is limited by their difficulty to be set apart from neural pogenitors, here we managed a good separation between these two by mapping human embryonic data onto a mouse reference data where tanycyte were clearly characterised. We found that comparing to RAX+ neural progenitors (SHH+, occur mostly at early embryonic stages), tanycytes express more radial glial genes FABP7, astrocyte marker AQP4, GFAP as well as tanycyte markers DIO2, CRYM, FRZB (Figure 2), in consistent with their radial glia-like identity as well as their function (Chinnaiya et al. 2022). Regarding human tanycyte heterogeneity, we identified similar tanycyte subtypes (α and β) to adult rodent tanycytes, though their gene expression profiles are slightly different that may reflect both species difference and/or functional maturity, for example, DIO2 and FGFR1 are more highly expressed in β tanycyte in adult mouse (Campbell et al. 2017), but in human embryonic tanycyte, they are similarly expressed in both α and β tanycytes (Figures 3 & S3). Another example, FABP7 is an α tanycyte marker in adult mouse (Campbell et al. 2017), however in human embryonic data, it is highly expressed in both embryonic tanycyte subtypes. In addition, GPR50 and FGF10 are expressed in both adult mouse tanycyte subtypes (Campbell et al. 2017), but absent/lowly expressed in our human embryonic tanycyte (Figure S3). The α and β tanycytes follow different developmental trajectories, but share the same origin and many dynamically upregulated genes (e.g. FABP7, CLU, CRYM). We anticipate our pioneer effect will facilitate more and more future studies on human tanycytes.

## Notes

### Competing Interest Statement

The authors have declared no competing interest.

