## Supplementary files for "Human embryonic tanycyte: heterogeneity and developmental trajectory"

**Supplementary Figure 1** Gather RAX$^+$ progenitors for downstream analysis. (A) UMAP plots of reprocessed RAX+ prognitor cells that were collected from different embryonic stages, showing the gene expression of *RAX* and *STMN2*. (B) UMAP plots of reprocessed RAX+ prognitor cells, colored based on the clusters acquired at Louvain clustering resolution 2.5. Newly emerged STMN2$^+$ and/or RAX$^{-/low}$ clusters "5", "8", "9", "12", "14", "15", "16", "18", "20", "23", "24", "29", "31", "33", "34", "35" were removed from further analyses. (C) New UMAP plot showing the cell cycle phases and embryonic stages of the remained RAX$^+$ STMN2$^{-/low}$ cells from step (B). Only non-cycling cells (phase G1) will be kept for downstream analysis. (D) UMAP showing the non-cycling RAX$^+$ STMN2$^{-/low}$ cells that will be used for downstream analyses.

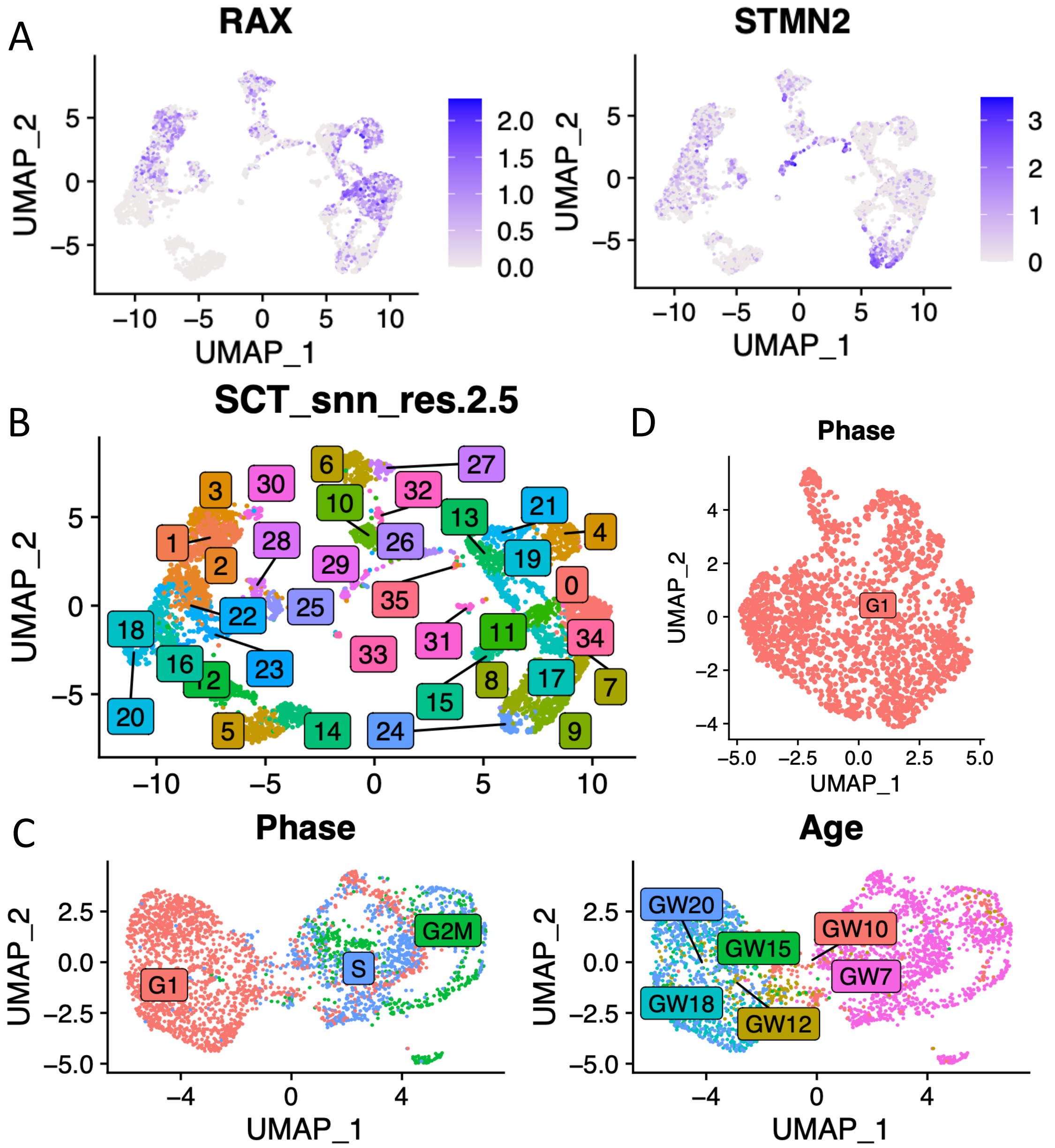

**Supplementary Figure 2.** UMAP plots showing the cell-wise expression levels of two Pars tuberalis marker genes (CCK and PITX2) as well as two tanycyte marker genes (CRYM and FRZB).

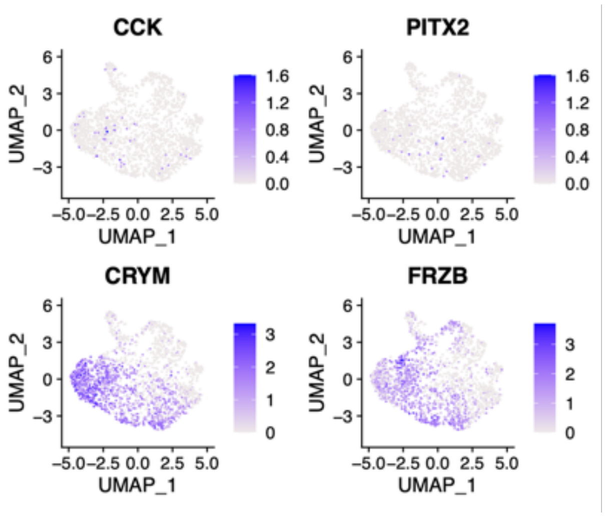


**Supplementary Figure 3** UMAP plots showing the cell-wise expression levels of three tanycyte marker genes.

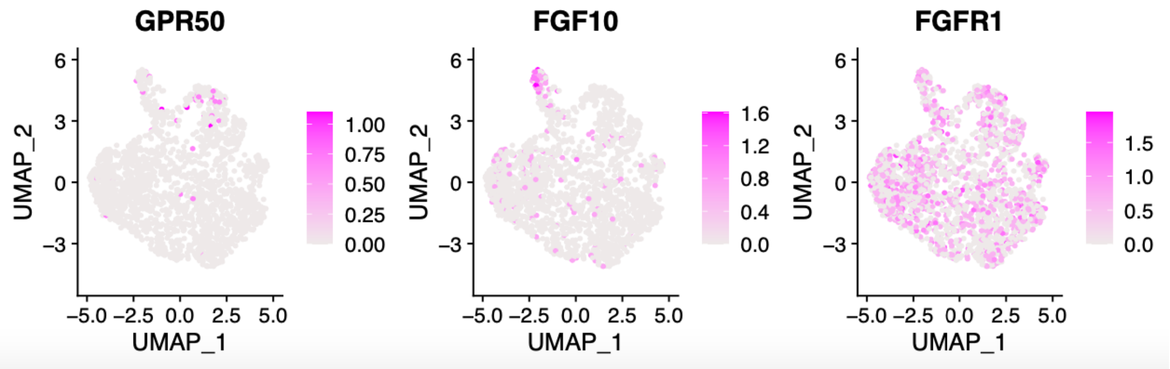

Supplementary Table 1 Predicted cell labels for each cluster at Louvain clustering resolution 1.5.

| **Louvain cluster (res=1.5)** | **Predicted label** |
|---|---|
| 0 | Tany |
| 1 | ParsTubers |
| 2 | Tany |
| 3 | Neuron |
| 4 | Neuron |
| 5 | Tany |
| 6 | Tany |
| 7 | Tany |
| 8 | Neuron |
| 9 | Tany |
| 10 | Neuron |
| 11 | Neuron |
| 12 | Neuron |
